# Longitudinal Exploration of Auditory Sensory-Perceptual Processing in CLN3 Disease (Juvenile Neuronal Ceroid Lipofuscinosis (Batten disease)): A High-Density Auditory Evoked Potential (AEP) Study

**DOI:** 10.1101/2025.11.19.689311

**Authors:** Erin K. Bojanek, Eve R. Lang, Heather R. Adams, Jennifer Vermilion, Erika F. Augustine, Tufikameni Brima, Shahzoda Nasimjonova, Edward G. Freedman, John J. Foxe

**Affiliations:** The Frederick J. and Marion A. Schindler Cognitive Neurophysiology Laboratory, Department of Neuroscience, University of Rochester School of Medicine and Dentistry, The Del Monte Institute for Neuroscience, Rochester, New York 14642, USA; The Golisano Intellectual and Developmental Disabilities Institute, Department of Neuroscience, University of Rochester School of Medicine and Dentistry, Rochester, New York 14642, USA; University of Rochester Batten Center (URBC), Department of Neurology and The Ernest J. Del Monte Institute for Neuroscience, University of Rochester School of Medicine and Dentistry, Rochester, New York 14642, USA; Kennedy Krieger Institute, Baltimore, Maryland 21205, USA

## Abstract

**Background:** There is currently limited information about sensory and perceptual abilities across the progression of CLN3 disease (Juvenile Neuronal Ceroid Lipofuscinosis; Batten disease), a recessively inherited lysosomal storage disorder and a leading cause of childhood neurodegeneration. Clinical symptoms include vision loss, motor impairments, and cognitive challenges, making it difficult to accurately assess neurocognitive and perceptual abilities. Thus, there is a critical need to identify objective biomarkers that can be used to inform disease progression and track treatment response in this population.

**Methods:** This exploratory study investigates longitudinal changes in auditory sensory perceptual processing in a small sample of individuals with genetically confirmed CLN3 disease (N=4; 3 male) compared to a cross-sectional sample of 60 neurotypical (NT) controls using high-density electroencephalography (EEG). We utilized a duration mismatch negativity (MMN) paradigm, identical to what has been used in our previous cross-sectional study. We examined the auditory evoked potentials (AEPs) of the standard tones across three different stimulus onset asynchrony conditions and examined the N1 and P2 components of the AEP.

**Results:** We found age related differences in the amplitudes of the N1 and P2 components in individuals with CLN3 disease relative to NT controls. These amplitude differences were most notable in CLN3 disease when participants were presented with standard tones that had the slowest presentation rate. Specifically, N1 and P2 amplitudes were more negative than NT controls in childhood and adolescence and as CLN3 disease participants aged, the amplitude of the AEPs was greater than controls. Further, a more positive N1 amplitude during the longest stimulus presentation condition was associated with both reduced verbal intelligence and working memory abilities in CLN3 disease participants.

**Conclusions:** Our preliminary findings parallel recently published work in a mouse model of CLN3 disease that showed both sex- and age-dependent disruptions in central auditory processing. Taken together, we demonstrate the utility of auditory EEG measures as a sensitive, objective and translational measure in CLN3 disease that may serve as a potential outcome measure useful in tracking disease progression. Continued work is needed in humans focused on sex-based differences and longitudinal changes of auditory processing in CLN3 disease.

## Introduction

Neuronal ceroid lipofuscinoses (NCL; Batten disease) are a family of rare, predominantly pediatric-onset, autosomal recessively inherited lysosomal storage disorders. Collectively, the NCLs represent the leading known cause of childhood neurodegenerative disorders (Boustany, 2013; Rider & Rider, 1999), with at least 13 distinct genetic forms of Batten disease. Juvenile neuronal ceroid lipofuscinosis (CLN3 disease) is one of the most common of these NCLs. CLN3 disease results from pathogenic variants in CLN3 that lead to pathological accumulation of ceroid lipofuscin in lysosomes of multiple cell types (Boustany, 2013; Munroe et al., 1997).

Classic CLN3 disease is characterized by onset of symptoms between 4-7 years of age followed by progressive neurologic decline with premature death typically in the third decade of life (Kuper et al., 2018; Masten et al., 2020; Mink et al., 2013). The first observed symptom is typically vision loss due to retinal degeneration that quickly progresses to blindness (Mink et al., 2013). Cognitive decline and learning difficulties are also often present around the time of diagnosis, though cognitive decline is not always recognized as early because changes in academic performance are frequently incorrectly attributed wholly to visual impairment (Kuper et al., 2018). Seizures frequently begin around 10-12 years of age (Augustine et al., 2015; Mink et al., 2013). Initial motor impairment is often reported around 10 years of age and early motor signs may include reduced walking speed (Adams et al., 2013; Kwon et al., 2011; Mink et al., 2013). Over time, parkinsonism (i.e., bradykinesia, impaired balance, and shuffling gait) and dysarthria emerge and progress over time. Impairments in cognitive ability eventually progress to dementia (Adams et al., 2007; Adams et al., 2013).

Given this complex set of symptoms and the progressive decline associated with CLN3 disease, it is challenging to accurately measure neurocognitive and perceptual abilities (Brima, Freedman, et al., 2024). Assessment of these abilities is further limited because the use of standard neurocognitive assessments may be neither appropriate nor feasible in this population given sensory impairment (vision loss), motor decline (loss of speech), and dementia. As such, there is currently limited knowledge about sensory and perceptual abilities across the full range of clinical stages of CLN3 disease, particularly in individuals at a more advanced disease state. Nevertheless, understanding sensory-perceptual abilities in CLN3 disease across disease progression is a necessary first step in identifying objective neurologic biomarkers (e.g., neuromarkers or endophenotypes) of CLN3 disease progression, which can be used to more accurately and objectively track disease progression and treatment response. Once identified, objective neuromarkers have the potential to serve as sensitive outcome measures that could be used to test the effectiveness of developing therapeutic approaches, and to aid in the objective staging of disease progression.

Current measures of CLN3 disease progression primarily rely on direct clinician observation and parent report, such as the Unified Batten Disease Rating Scale (UBDRS), which measures disease progression across key symptom domains (i.e., physical, seizure, behavior, and capability; (Marshall et al., 2005). This scale has proven beneficial in characterizing the natural history of CLN3 disease over the last 20 plus years (Kwon et al., 2011; Marshall et al., 2005). Distilled from elements of the UBDRS, the CLN3 Staging System (CLN3SS) was developed to classify affected individuals into one of four discrete disease stages (Masten et al., 2020). The CLN3SS applies clinically meaningful features from the UBDRS to classify individuals into 4 distinct, incremental stages (stages 0-3), with higher CLN3SS denoting more advanced disease: (0) genetic confirmation, (1) onset of vision loss, (2) occurrence of one or more seizures, and (3) inability to walk without assistance (Masten et al., 2020). Each higher stage requires the presence of the clinical feature from each preceding stage. The UBDRS and CLN3SS have been beneficial for stratifying patients into subgroups, yet these scales are limited in their sensitivity to detect change over a short period of time, particularly given that CLN3 disease has a slowly progressing disease course that unfolds over two or more decades. Thus these measures may be limited in their use as outcome measures for short-term clinical trials (Masten et al., 2020).

Clinical trials in rare disease populations have faced a variety of challenges including small sample sizes, lack of translatable outcome measures between animal models and human trials, and outcome measures that are not sensitive to changes during a short term trial (Goodspeed et al., 2023). Thus, there is considerable need for objective biomarkers of brain differences. One readily deployable method for the identification of objective neuromarkers is event-related potential (ERP) recordings using electroencephalography (EEG). ERPs are an appealing translational method in rare disease research as they can be used to measure brain activity in both humans and animal models (Brima et al., 2019; Ding et al., 2025; Foxe et al., 2016; Francisco et al., 2020; LeBlanc et al., 2015; Sysoeva et al., 2020). EEG allows for tracking of the transmission of sensory information with high temporal precision and provides a measure of sensory processing to specific sensory stimuli, not readily available with other non-invasive methods. ERPs can be measured in the absence of an overt behavioral response, which is particularly advantageous when disability impedes motor and verbal responses (Mills et al., 2004; Riva et al., 2018; Yoder et al., 2006). EEG can also provide a measure of information flow from early sensory perception to higher level cognitive stages of processing (Foxe & Simpson, 2002; Näätänen, 2003), and they show a high level of test-retest reliability, making them an ideal measure for use in longitudinal studies and clinical trials (Beker et al., 2021; Kileny & Kripal, 1987; Malcolm et al., 2019). While there is currently limited research using EEG recordings in CLN3 disease, there has been a wide range of research examining EEG and ERP recordings in a range of neurodevelopmental disorders including autism (Brandwein et al., 2015; Jeste & Nelson III, 2009; Russo et al., 2010; Yuan et al., 2025) and rare disease populations (Brima, Beker, et al., 2024; Francisco et al., 2020; Goodspeed et al., 2023). There is emerging evidence of quantitative EEG analysis methods (e.g., spectral power analyses, measures of connectivity, ERP analyses) being used to identify potential objective biomarkers of disease severity in Angelman syndrome (Martinez et al., 2023; McDougle & Keary, 2019), MECP2 disorders (Brima, Beker, et al., 2024; Brima et al., 2019; Foxe et al., 2016; Sysoeva et al., 2020) and Fragile X syndrome (Hooper et al., 2021; Wang et al., 2017; Wilkinson & Nelson, 2021). This emerging research suggests that quantitative EEG analysis methods should be explored as a helpful tool in the identification of candidate biomarkers in other rare genetic neurodevelopmental disorders (Goodspeed et al., 2023).

Given the early central vision loss in individuals with CLN3 disease, we chose to measure auditory sensory-perceptual processing as a direct assay of the integrity of early cortical processing. This study therefore focuses on the measurement of basic auditory processing, which is foundational for later levels of auditory processing and language understanding. Our group has previously examined auditory discrimination and auditory sensory memory in a cross-sectional study of participants with CLN3 disease (Brima, Freedman, et al., 2024) using the well-characterized mismatch negativity (MMN) component of the ERP (Brima, Beker, et al., 2024; Näätänen, 2003; Ritter et al., 2002). Brima et al. found that individuals with CLN3 disease show a partially preserved ability to automatically detect duration deviations when stimuli are presented at a medium presentation rate (circa 1 Hz), yet they showed deficits in their ability to detect changes when the presentation of the auditory stimuli was slower (∼0.5 Hz) or faster (∼2 Hz). These findings suggest that individuals with CLN3 disease have some preserved auditory perceptual abilities, yet when additional demands are put on the auditory system, they show clear deficits which likely have implications for information processing and language understanding (Brima, Freedman, et al., 2024). However, research examining neural mechanisms and integrity of central auditory processing abilities has been limited, particularly examination of auditory processing longitudinally and across disease progression in individuals with CLN3 disease.

This exploratory study characterizes basic auditory processing in a sample of four individuals with CLN3 disease across disease stages. We used objective, noninvasive EEG to examine auditory evoked potentials (AEPs) over multiple time points and compared these neurophysiological measures to clinical measures of disease severity and cognition. The primary aim of this exploratory study is to provide early evidence of the utility of EEG measures as an objective outcome measure that may be beneficial in tracking disease progression over time. The AEPs of CLN3 disease participants were also compared to a large normative cross-sectional sample of neurotypical (NT) control participants (N = 60; child, adolescent, and adult) who completed the same EEG tasks at a single timepoint. The modest sample size of CLN3 disease participants in this study is a known limitation of rare disease work, yet the advantage in the present study was the ability to longitudinally evaluate AEP changes in participants with CLN3 disease.

## Methods

### Participants

Four participants with genetically confirmed CLN3 disease and 60 NT controls were included in this mixed longitudinal and cross-sectional study. CLN3 disease participants were recruited through the University of Rochester Batten Center and NT controls were recruited from the community. Participants with CLN3 disease (3 male) repeated an EEG task at multiple timepoints (4-6), each at least 6 months apart. Participants with CLN3 disease were between the ages of 11-19 years old at their first visit and 15-25 years old at their final visit. Participants with CLN3 disease were primarily seen annually at the Batten Disease Support, Research, and Advocacy (BDSRA) Foundation annual family conference, when they attended, in addition to visits to the University of Rochester.

Participants in the NT control group (ages 7-30 years) participated in this study at only one timepoint. The NT control group was split into three age-bands (e.g., child, adolescent, adult) to more closely match the ages of the CLN3 disease participants and to group NT controls by developmental trajectory of the AEP (Bishop et al., 2011; Bishop et al., 2007; Ponton et al., 2000). NT control participants ages 7 to 12 years were in the child control group (N=19) and served as comparisons for CLN3 disease participants at ages 11 and 12 years. NT control participants aged 13 to 18 years were in the adolescent control group (N=19) and served as comparisons for CLN3 disease participants ages 13 to 18 years. NT control participants aged 19 to 30 were in the adult control group (N=22) and served as comparisons for CLN3 disease participants ages 19 to 25 years (NT control age bands are described in more detail in Table 1). This method allowed us to compare AEPs from CLN3 disease participants across various ages to NT control participants that were close in age given developmental changes that occur to the AEP (Brandwein et al., 2011; Gomes et al., 2001; Gomes et al., 1999).

**Table 1.** Demographic characteristics of NT control participants.

| **Group** | **N (# Male)** | **Age Range** | **Mean Age (SD)** |
| --- | --- | --- | --- |
| Child | 19 (7) | 7-12 | 8.9 (1.9) |
| Adolescent | 19 (8) | 13-18 | 15.5 (1.4) |
| Adult | 22 (11) | 19-30 | 22.6 (3.2) |

Participants with CLN3 disease were not included in this study if they met any of the following criteria: onset of seizures before age 4 years or developmental concerns not related to CLN3 disease prior to age 4 years. Participants in both groups were also not included if they had uncorrected hearing loss or an ear infection on the day of EEG acquisition. NT control participants were excluded if they had a neurodevelopmental disorder or a neurodevelopmental disorder in a first-degree biological family member, or any self-reported or parent-reported neurological or psychiatric disorders.

Participants with CLN3 disease completed detailed clinical phenotyping and their parents provided detailed medical history. All four participants had genetically confirmed CLN3 disease (Masten et al., 2021). Symptom severity of CLN3 disease was measured using the UBDRS (Marshall et al., 2005), administered by a trained clinician, and disease stage was assigned using the CLN3SS (Masten et al., 2020). There was little change in participants CLN3SS across visits (Table 2). Clinical phenotyping for CLN3 disease participants included verbal components of the Wechsler Intelligence Scale for Children, Fifth Edition (WISC-V) for all participants. Given the cognitive decline present in older individuals with CLN3 disease, for participants 17 years or older, the ceiling age band norms (age 16 years, 11 months) were used to generate scaled and standard scores. We calculated participants’ verbal comprehension index (VCI) standard score (mean=100, SD = 15), and the digit span forward (DSF) scaled score (mean=10, SD=3). There were several timepoints at which participants could not complete cognitive testing due to interfering behaviors (i.e. Sub #2 time point 1), or due to advanced disease stage including limited verbal abilities (i.e., Sub #3 & Sub #4). Cognitive testing showed that Sub #1 retained average skills throughout the study, with only a slight drop overall, Sub #2 retained below average skills, and Sub 3 & 4 showed cognitive skills near the floor across assessments. Demographic information and clinical characterization of CLN3 disease participants across time points is reported in more detail in Table 2. Clinical information from the UBDRS participants including age at symptom onset are reported in Table 3.

**Table 2.** Demographic and clinical characteristics of CLN3 disease participants.

| **Subject** | **Timepoint** | **Age** | **CLN3SS** | **VCI**<br>**Standard**<br>**Score** | **DSF**<br>**Scaled**<br>**score** |
| --- | --- | --- | --- | --- | --- |
| Sub #1 (male) | 1 | 12 | 1 | 96 | 10 |
|  | 2 | 14 | 2 | 86 | 10 |
|  | 3 | 15 | 2 | 89 | 9 |
|  | 4 | 18 | 2 | 89 | 10 |
| Sub #2 (female) | 1 | 11 | 2 | -- | -- |
|  | 2 | 12 | 2 | 78 | 5 |
|  | 3 | 12 | 2 | 84 | 3 |
|  | 4 | 13 | 2 | 76 | 2 |
|  | 5 | 15 | 2 | 73 | 2 |
| Sub #3 (male) | 1 | 19 | 2 | 47 | 3 |
|  | 2 | 20 | 2 | 45 | 1 |
|  | 3 | 21 | 2 | 45 | 1 |
|  | 4 | 21 | 2 | 45 | 1 |
|  | 5 | 25 | 3 | -- | -- |
| Sub #4 (male) | 1 | 19 | 2 | 59 | 2 |
|  | 2 | 20 | 3 | 50 | 2 |
|  | 3 | 20 | 3 | 55 | 1 |
|  | 4 | 21 | 3 | 55 | -- |
|  | 5 | 22 | 3 | -- | -- |
|  | 6 | 25 | 3 | -- | -- |

Note. CLN3SS = CLN3 Staging System, VCI = Verbal Comprehension Index standard score, and DSF=Digit Span Forward scaled score at each timepoint. "--" denotes data not collected because participant was non-compliant or unable to complete task.

**Table 3.** Age of onset of clinical symptoms for CLN3 disease participants.

|  | **Vision**<br>**Loss** | **Sleep** | **Cognitive** | **Behavioral** | **Seizure** | **Motor** | **Weight**<br>**Loss/Feeding** |
| --- | --- | --- | --- | --- | --- | --- | --- |
| Sub #1 | 9 | 10 | 11 | 11 | 13 | n/a | n/a |
| Sub #2 | 7 | n/a | 7 | 8 | 10 | n/a | n/a |
| Sub #3 | 5 | 6 | 5 | 9 | 13 | 15 | 20 |
| Sub #4 | 5 | 6 | 7 | 7 | 7 | 14 | n/a |

Note. Onset of clinical symptoms of CLN3 disease recorded from Unified Batten Disease Rating Scale (UBDRS) for each participant.

### Experimental Design

Participants were presented with an auditory MMN paradigm while recording EEG. Experimental procedures are identical to those described in our prior work (Brima, Freedman, et al., 2024; Foxe et al., 2016). Middle ear conductive hearing loss was ruled out in all participants via tympanometry. NT control participants sat in a sound attenuated and electrically shielded booth (Industrial Acoustics Company, Bronx, NY). Recording from CLN3 disease participants took place either in the same electrically shielded booth or in an offsite, private setup at the BDSRA Foundation annual family conference. CLN3 disease participants either sat in a chair or their wheelchair, depending on their preference. All participants were offered fidget toys or NT control participants watched a silent video while passively listening to auditory stimuli presented at 75 dB SPL using headphones (Sennheiser electronic GmbH & Co. KG, USA). The task was created using Presentation Software (Neurobehavioral Systems, Inc. Berkeley, CA). An oddball paradigm was used, standard tones occurred 85% of the time and deviant tones occurred 15% of the time. Standard tones were 100ms in duration, 1000Hz, and had a rise and fall time of 10ms. Deviant tones were 180ms in duration but otherwise identical. Tones were presented at different stimulus onset asynchronies (SOAs). The SOAs were 450, 900, or 1800 ms. Each block consisted of only one SOA. Blocks were fixed in length so there are a different number of stimuli per block depending on SOA (i.e. 450ms = 500 trials/block, 900ms = 250trials/block, 1800ms = 125trials/block). The order of the blocks was randomized, and participants completed 14 blocks (i.e., 2×450 SOA, 4×900 SOA, 8×1800 SOA) to total 1000 trials per condition lasting about 70 minutes. The present study analyzed only electrophysiological responses (AEP) to the standard 100ms tones.

### EEG Acquisition

A BioSemi ActiveTwo system (BioSemi B.V. Amsterdam, Netherlands) with either a 32 or 64 channel electrode array was used to collect EEG data. Data from 1 NT child, 1 NT adolescent, and 7 NT adults were recorded using a 32-channel cap. Additionally, for 2 CLN3 disease participants, the 32-channel cap was used at 3 timepoints, and for the remaining 2 CLN3 disease participants, a 32-channel cap was used at 4 timepoints. All other NT recordings and additional CLN3 disease participant timepoints utilized a 64-channel cap. Electrodes were positioned using the BioSemi Equiradial system, with 2 external electrodes over the mastoids and one on the outside of each eye. EEG data were referenced online to an active and passive electrode, common mode sense (CMS), and driven right leg (DRL), respectively. Analyses focused specifically on the N1 and P2 components of the AEP. The N1 component is associated with initial orientation to sound, while the P2 component indicates further auditory processing and learning. The windows of measurement were defined as the peak within 80 to 130 ms for the N1, while the P2 component was defined as the peak within 130 to 190 ms.

### EEG Data Processing

EEG data were analyzed offline using custom scripts that included EEGLAB toolbox (Delorme & Makeig, 2004) and Fieldtrip toolbox (Oostenveld et al., 2011) functions for MATLAB (version 2024b; MathWorks, Natick, MA, USA). Data were filtered using a Chebyshev II spectral filter with a bandpass of 1-40 Hz. A channel rejection algorithm, which identified bad channels using measures of standard deviation and covariance with neighboring channels (6 channels) was utilized to identify bad channels. Rejected channels were then replaced through spherical spline interpolation (EEGLAB). Data were then re-referenced to the average reference. Data were epoched from −100ms to 400ms. Epochs were rejected if the maximum value was >150 uV or >2 standard deviations away from the mean max value in either direction. The mean number of accepted trials and mean number of rejected channels for each condition and group is reported in Supplemental Table 1.

Composite averages were generated from F3, Fz, and F4 scalp electrodes when the 32-channel cap was used and from FC3, FCz, and FC4 when the 64-channel cap was used. Prior literature shows a strong presence of the AEP at both sets of scalp locations, and highlights that these scalp locations can be used interchangeably (Brima, Freedman, et al., 2024).

### Statistical Analyses

To provide normal distributions of NT control data against which we could compare the CLN3 disease participant data, we calculated the grand average and standard deviations of the peak amplitude and latency of the N1 and P2 components separately for each of the NT control groups. To examine neurotypical age-related changes in the AEP, we ran a repeated measures ANOVA for the NT control participants with SOA condition (i.e., 450 ms, 900 ms, 1800 ms) as the within subject factor and group (i.e., NT child, NT adolescent, and NT adult) as the between-subject factor. Differences between AEPs of CLN3 disease participants at each time point and the neurotypical distribution were examined using Z-scores, which were calculated for each CLN3 disease participant at each time point using the age-matched NT control group (e.g., child, adolescent, or adult) as the sample mean and sample standard deviation. A negative z-score denotes a reduced amplitude compared to the control group and a positive z-score denotes an increased amplitude compared to the age-matched control comparison.

Associations between the amplitudes of the N1 and P2 components across SOA conditions and clinical measures (CLN3SS, VCI Standard Score, and DSF Scaled Score) were examined using Spearman correlations for the CLN3 disease participants. We also examined the association between age and N1 and P2 component amplitudes across SOA conditions separately for the CLN3 disease and NT control participants using Pearson correlations. Given the small number of participants in the CLN3 cohort, only correlations |r|>0.5 and p<0.01 were interpreted as significant.

## Results

### EEG Results

#### Control Participants

We examined the amplitude of the N1 and P2 components of the auditory evoked potential (AEP) in response to standard tones across the three SOA conditions in participants with CLN3 disease and NT control participants. In the NT control participants, we found a significant SOA x age-group interaction (F(2.630, 57)=9.331, p<0.001, *η*^2^ =0.247) in which the N1 amplitude increased in negativity with age and across SOA conditions, with the strongest negativity seen in the adult control participants during the 1800 ms SOA condition. For the P2 amplitude, we did not find a significant SOA x age-group interaction (p=0.128, *η*^2^ =0.062). Mean N1 and P2 amplitudes for NT control participants across SOA are reported in Tables 4 and 5, respectively.

**Table 4.** N1 Amplitudes for NT control participants.

|  | NT Child | NT Adolescent | NT Adult |
| --- | --- | --- | --- |
| 450ms SOA | 0.720 (0.812) | 0.099 (0.457) | -0.106 (0.468) |
| 900 ms SOA | 0.583 (1.028) | -0.535 (0.781) | -1.242 (0.735) |
| 1800 ms SOA | -0.172 (1.184) | -1.264 (1.255) | -2.252 (0.978) |

Note. Data are reported as mean (standard deviation).

**Table 5.** P2 Amplitudes for NT control participants.

|  | NT Child | NT Adolescent | NT Adult |
| --- | --- | --- | --- |
| 450 ms SOA | 0.975 (0.783) | 0.605 (0.485) | 0.373 (0.519) |
| 900 ms SOA | 1.254 (1.082) | 0.598 (0.548) | 0.245 (0.892) |
| 1800 ms SOA | 1.036 (1.165) | 0.822 (0.706) | 0.343 (1.015) |

Note. Data are reported as mean (standard deviation).

#### CLN3 Disease Participants

When examining z-scores of CLN3 disease participants compared to the normal distribution of age matched NT controls, we found that younger CLN3 disease participants (Sub #1 & 2), showed a more negative N1 amplitude relative to the NT child group, particularly at the longer SOA conditions (Figure 1A). Adolescent CLN3 disease participants (Sub 1 &2) compared to the NT adolescent group showed a similar N1 amplitude during the 450 ms SOA condition but a slightly more negative amplitude compared to the NT adolescent group during the 900 ms and 1800 ms SOA conditions. By adulthood, CLN3 disease participants (Sub 3 &4) showed N1 amplitudes similar to the NT adult group at the 450 ms SOA condition and more positive N1 amplitudes during the 900 ms and 1800 ms SOA conditions. This pattern of increased N1 negativity in younger CLN3 disease participants and reduced negativity in older CLN3 disease participants is opposite to that observed in NT control participants across development.

**Figure 1.**
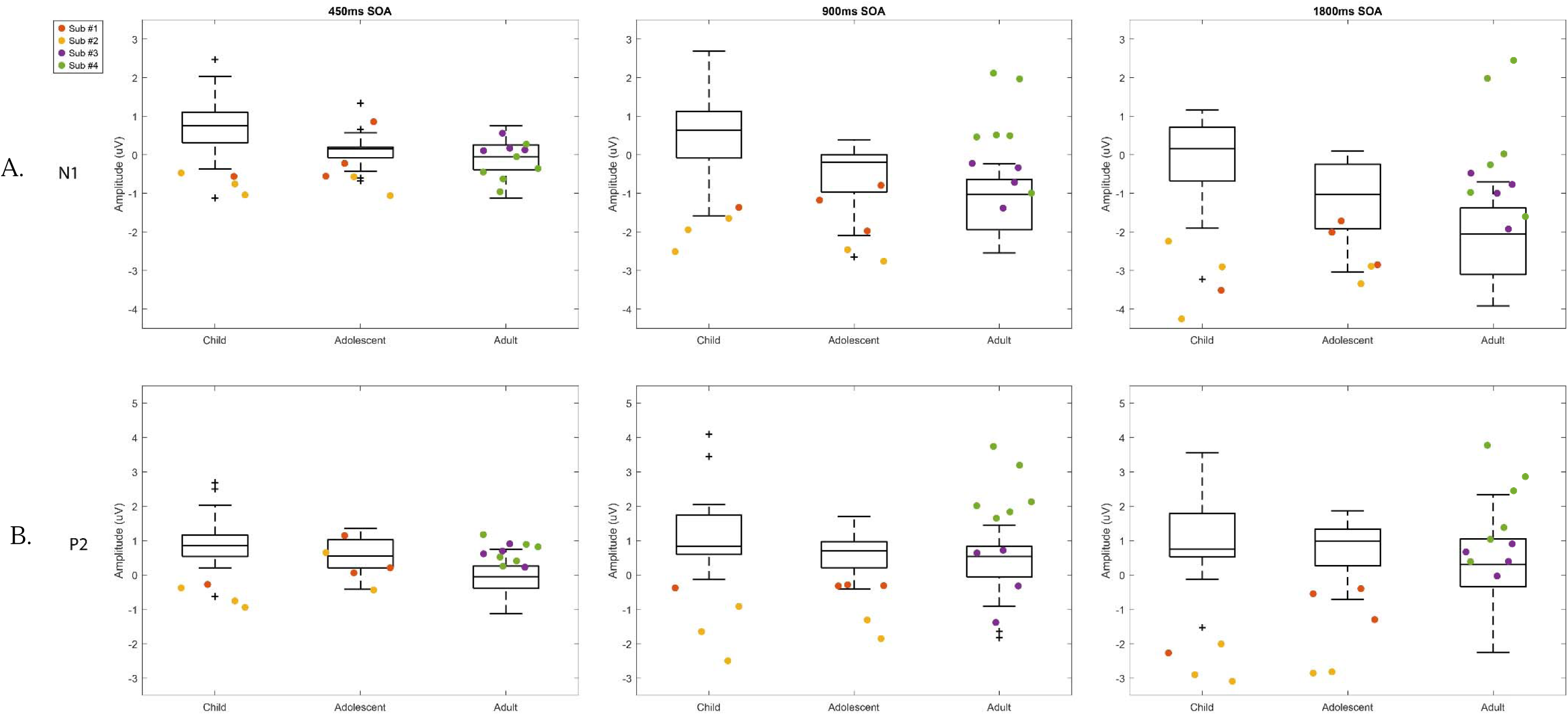
Boxplots of mean AEP mean amplitudes for N1(A) and P2 (B) components across SOA conditions. NT controls are shown in the box and whisker plots in black with CLN3 disease participants shown in color. CLN3 disease and NT control participants show opposite patterns of AEP response across development in both N1 and P2 AEP components.

We found a similar pattern in the CLN3 disease participants in their P2 amplitudes as their N1 amplitudes. Younger CLN3 disease participants, particularly at the longer SOA conditions, showed a less positive P2 amplitude relative to the NT child group (Figure 1B). Similar to the N1 amplitude, by adulthood, CLN3 disease participants showed more positive P2 amplitudes than the NT adult group, particularly at longer SOA conditions.

We also examined the amplitudes of ERPs separately for each SOA condition and each CLN3 disease participant individually across timepoints (Figure 2). Subjects 1 and 2 were ages 12 and 11 years old respectively at their baseline assessments. Compared to the normal distribution of ERP amplitudes in NT controls, subject 1, had the greatest differences (0.4 to 2.8 standard deviations difference) in the longer SOA conditions. However, across timepoints, the N1 (Table 6) and P2 (Table 7) amplitudes of subject 1 were closer to the normal distribution of amplitudes seen in NT control participants. In subject 2, the P2 component, particularly in the longer SOA conditions, differed most compared to NT child and adolescent control groups (2.6 to 5.2 standard deviations difference). There was no consistent pattern of change across timepoints in subject 2 as seen in subject 1.

**Figure 2.**
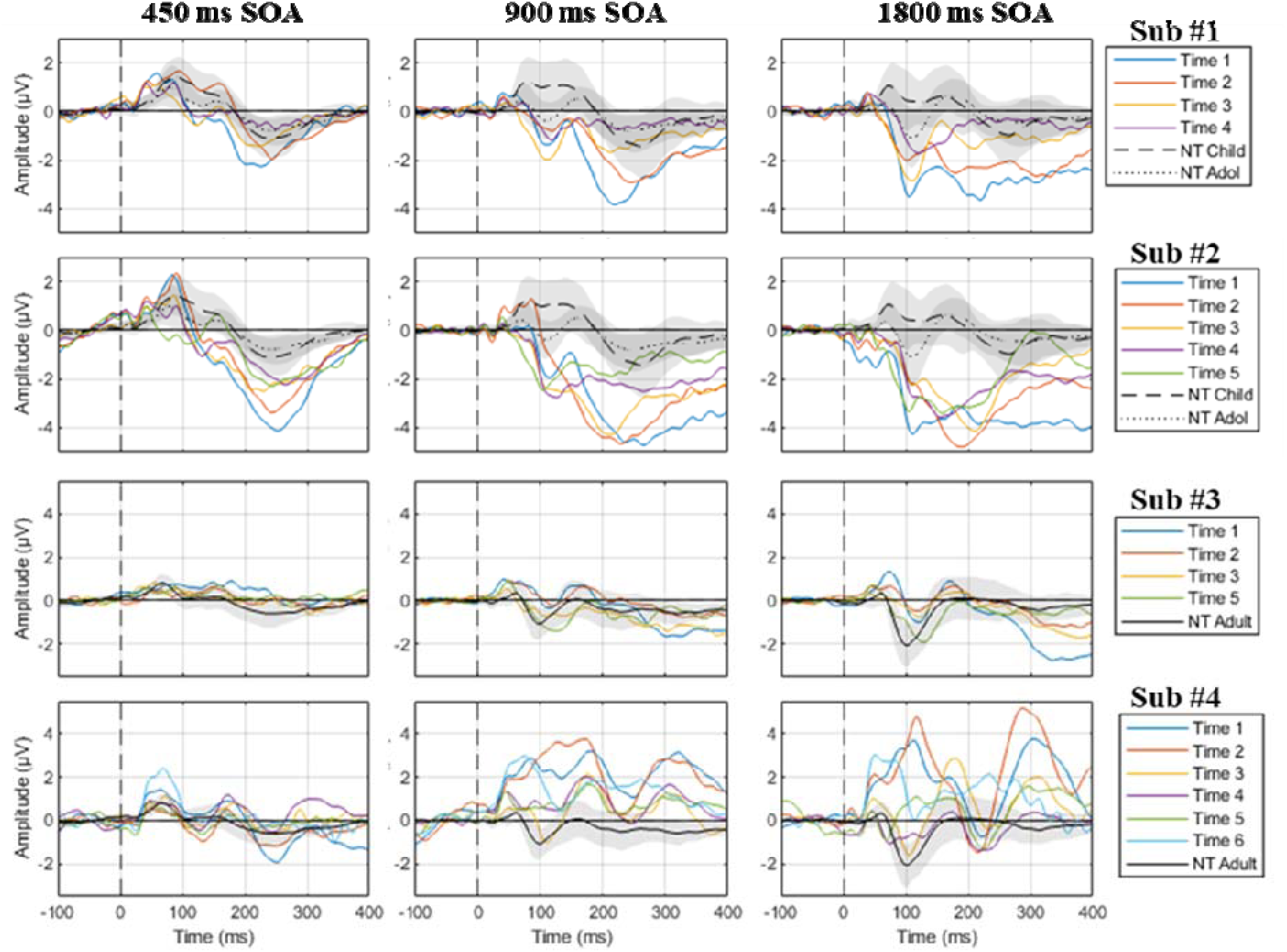
ERPs of each CLN3 disease participant separately across each time point and SOA. The aged-matched NT control group is included in black for each participant. The shaded regions denote the standard deviation of the corresponding NT control group. Sub #3, Time 4 is excluded from this figure due to a significant seizure less than 24 hours before research participation that required use of rescue medication. Data for this timepoint is included in Supplemental Figure 1 due to abnormal seizure activity and medication use.

**Table 6.**
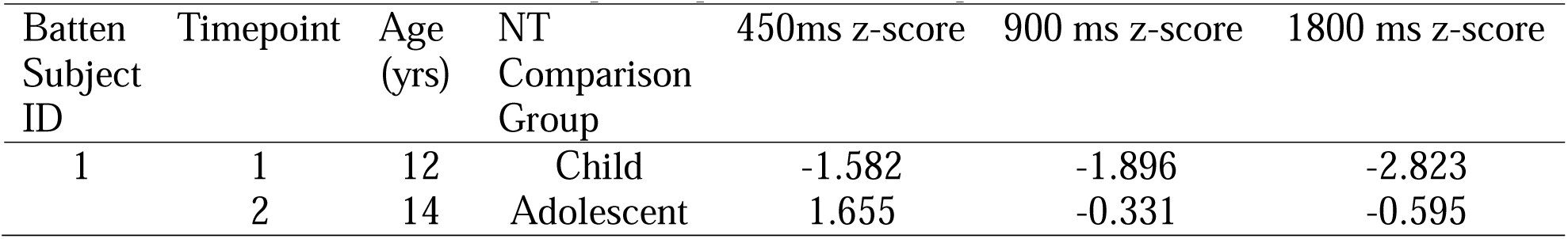
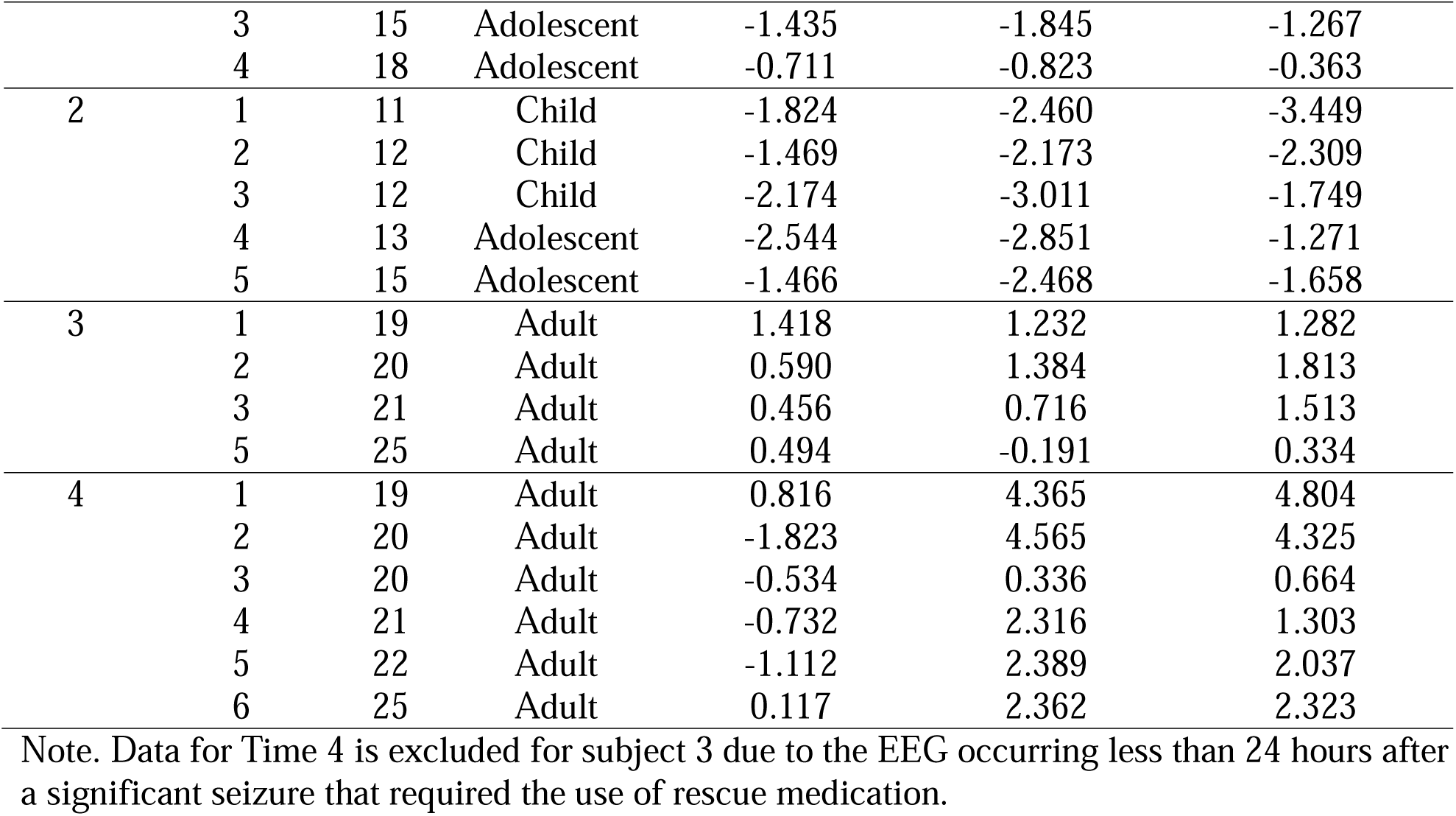
Z-scores for CLN3 disease participants for N1 Amplitudes across SOA conditions.

**Table 7.**
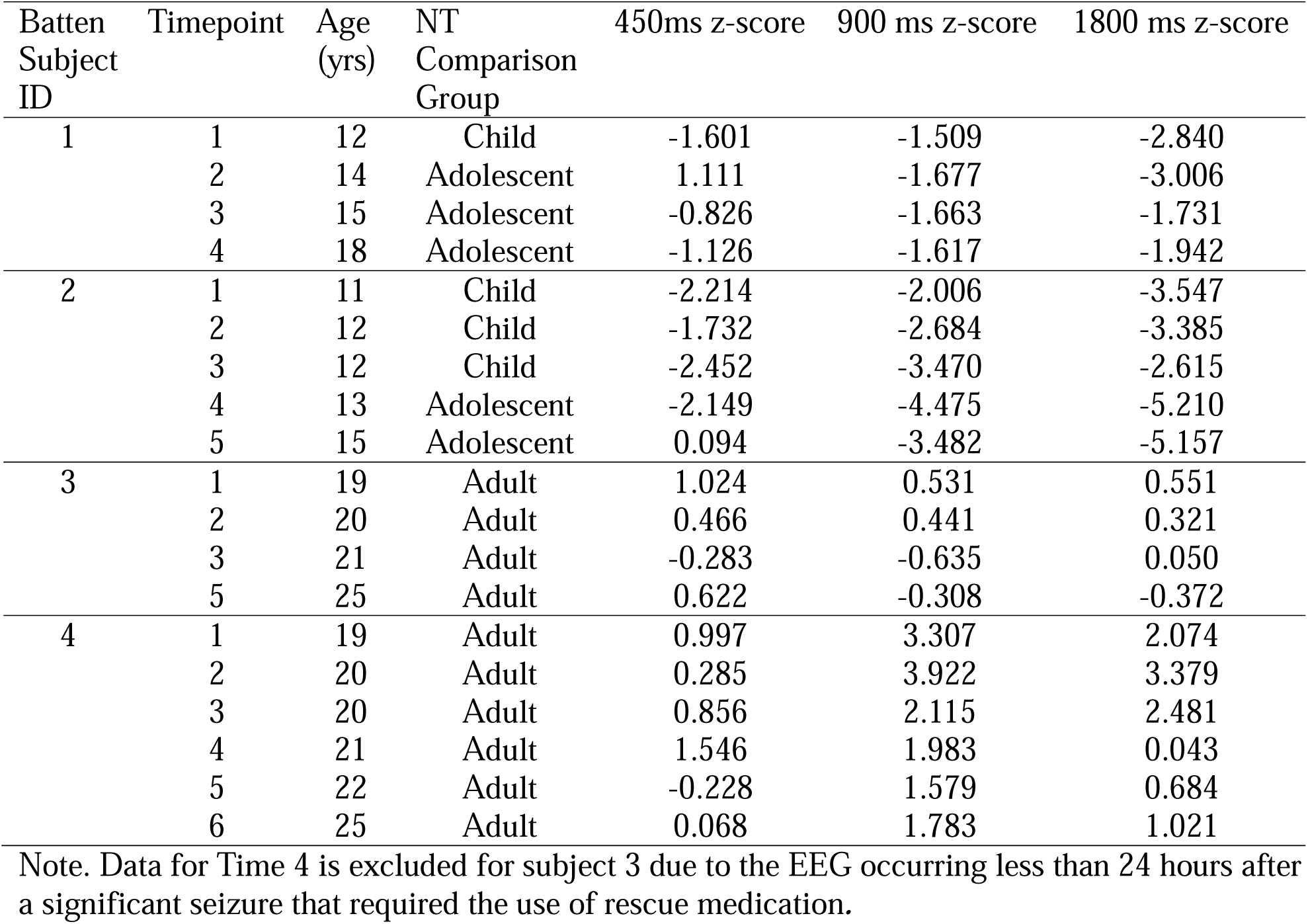
Z-scores for CLN3 disease participants for P2 Amplitudes across SOA conditions.

| Batten Subject ID | Timepoint | Age (yrs) | NT Comparison Group | 450ms z-score | 900 ms z-score | 1800 ms z-score |
| --- | --- | --- | --- | --- | --- | --- |
| 1 | 1 | 12 | Child | -1.601 | -1.509 | -2.840 |
|  | 2 | 14 | Adolescent | 1.111 | -1.677 | -3.006 |
|  | 3 | 15 | Adolescent | -0.826 | -1.663 | -1.731 |
|  | 4 | 18 | Adolescent | -1.126 | -1.617 | -1.942 |
| 2 | 1 | 11 | Child | -2.214 | -2.006 | -3.547 |
|  | 2 | 12 | Child | -1.732 | -2.684 | -3.385 |
|  | 3 | 12 | Child | -2.452 | -3.470 | -2.615 |
|  | 4 | 13 | Adolescent | -2.149 | -4.475 | -5.210 |
|  | 5 | 15 | Adolescent | 0.094 | -3.482 | -5.157 |
| 3 | 1 | 19 | Adult | 1.024 | 0.531 | 0.551 |
|  | 2 | 20 | Adult | 0.466 | 0.441 | 0.321 |
|  | 3 | 21 | Adult | -0.283 | -0.635 | 0.050 |
|  | 5 | 25 | Adult | 0.622 | -0.308 | -0.372 |
| 4 | 1 | 19 | Adult | 0.997 | 3.307 | 2.074 |
|  | 2 | 20 | Adult | 0.285 | 3.922 | 3.379 |
|  | 3 | 20 | Adult | 0.856 | 2.115 | 2.481 |
|  | 4 | 21 | Adult | 1.546 | 1.983 | 0.043 |
|  | 5 | 22 | Adult | -0.228 | 1.579 | 0.684 |
|  | 6 | 25 | Adult | 0.068 | 1.783 | 1.021 |

Note. Data for Time 4 is excluded for subject 3 due to the EEG occurring less than 24 hours after a significant seizure that required the use of rescue medication.

Subjects 3 and 4 were both ages 19 to 25 years old across 5 and 6 timepoints respectively. Subject 3 showed amplitudes of N1 (Table 6) and P2 (Table 7) components that were closest to NT adult comparison amplitudes at older ages. Amplitudes for the N1 component were less than 2 standard deviations different from the NT adult group for the N1 component and less than about 1 standard deviation difference for the P2 component compared to the NT adult group. In subject 3, there was one outlier at timepoint 4, this visit occurred less than 24 hours after the participant had a significant seizure that required the use of rescue medication. Therefore, data for this visit is included in Supplemental Figure 1 but is not included in the primary figures or tables due to the abnormal seizure activity and medication use. Subject 4 showed abnormal N1 and P2 amplitudes relative to the NT adult group. Subject 4’s N1 amplitude was less negative than the NT adult group across the longer SOA conditions (900 and 1800 ms), N1 amplitudes were closest to the NT adult group performance at timepoint 3 when subject 4 was at 20 years old (less than 1 standard deviation difference), however, the amplitudes were more positive again at later timepoints (1.3 to 2.4 standard deviations difference). P2 amplitudes were more positive than the NT adult group across the longer SOA conditions (900 and 1800 ms) across all timepoints.

#### Clinical Correlations

We examined the association between AEPs and clinical symptoms of CLN3 disease (Supplemental Table 2). CLN3 disease participants showed a significant positive association between the CLN3SS and P2 amplitude at 900 ms (r=0.650, p=0.003; Figure 3A) and 1800 ms (r=0.607, p=0.006; Figure 3B). There was also a significant negative association between the N1 amplitude at 1800 ms and the verbal comprehension index (r=-0.713, p=0.003; Figure 4A) and the digit span forward scaled score (r=-0.675, p=0.006; Figure 4B).

**Figure 3.**
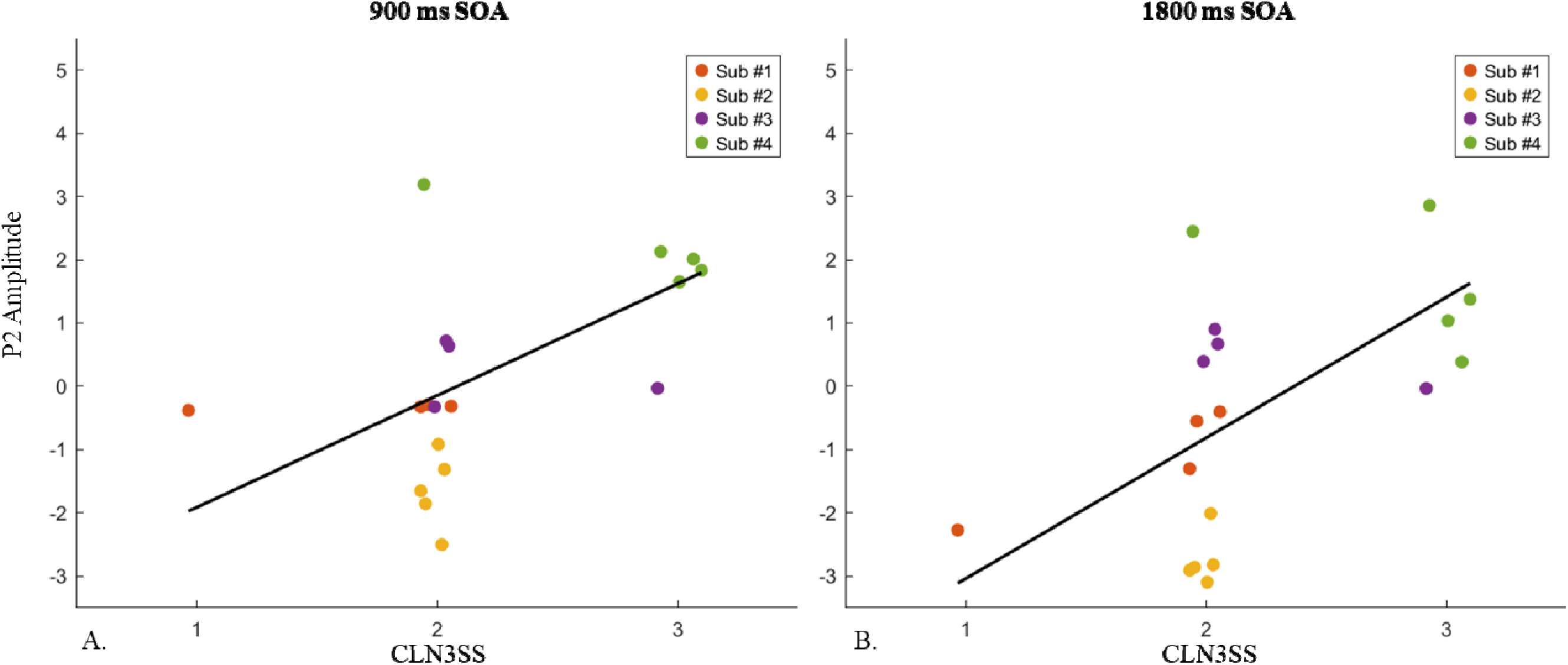
Significant associations between P2 amplitude in CLN3 disease participants and clinical symptoms of CLN3 disease as measured by the CLN3SS for the 900 ms SOA condition (A) and 1800 ms SOA condition (B). There was a significant positive association between CLN3SS and P2 amplitude at 900 ms and CLN3SS (r=0.650, p=0.003; A) and 1800 ms (r=0.607, p=0.006; B).

**Figure 4.**
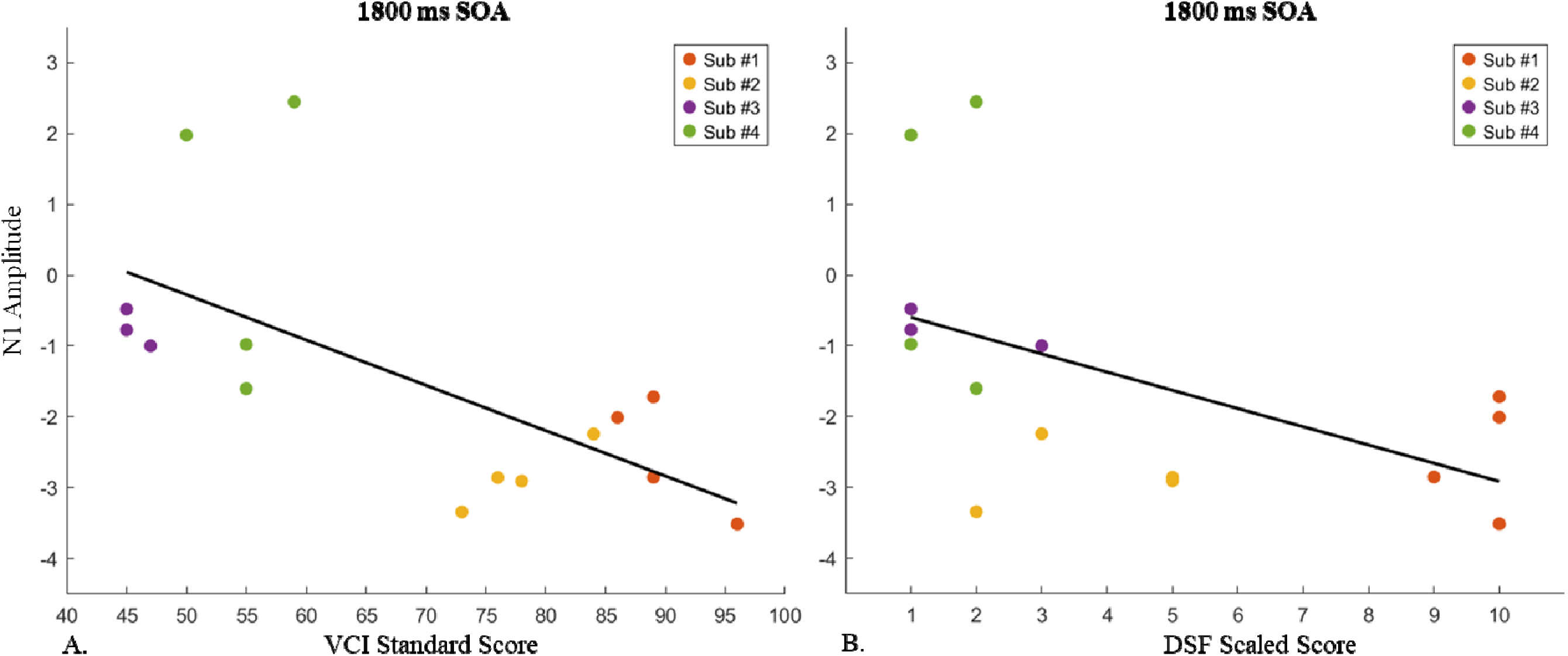
Significant clinical associations between N1 amplitude in CLN3 disease participants and Verbal Comprehension Index (VCI) standard score (A) and Digit Span Forward (DSF) Scaled Score (B) for the 1800 ms SOA conditions. There was a significant negative association between the N1 amplitude at 1800 ms and the VCI (r=-0.713, p=0.003; A) and the DSF scaled score (r=-0.675, p=0.006; B).

We also examined the association of AEPs and age in CLN3 disease and NT control participants. CLN3 disease participants showed a significant positive association between age and N1 amplitude at 900 ms (r=0.615, p=0.005) and 1800 ms (r=0.656, p=0.002) as well as between age and P2 amplitude at 450 ms (r=0.620, p=0.005), 900 ms (r=0.671, p=0.002), and 1800 ms (r=0.756, r<0.001). NT control participants showed a significant negative association between age and N1 amplitude at 900 ms (r=-0.613, p<0.001) and 1800 ms (r=-0.568, p<0.001). Additional results of age associations are reported in Supplemental Table 3 and Supplemental Figure 2.

## Discussion

In this study, we found that using EEG to quantify AEPs is a novel approach to assess basic auditory processing longitudinally in CLN3 disease. By following patients longitudinally, we have shown that there are emerging age-related differences across disease stages in individuals with CLN3 disease compared to age-matched NT controls. While this work is limited by the modest sample size and preliminary nature, it highlights the promise of these objective EEG methods in quantifying central auditory processing differences as a possible objective measure, useful as a clinical outcome measure and to track disease progression.

This current work builds on previous literature examining auditory processing in individuals with CLN3 disease using the same MMN paradigm (Brima, Freedman, et al., 2024). Using a cross-sectional design, Brima et al., showed that individuals with CLN3 disease (N=21, age range 6-28 years) maintained the ability to automatically detect and decode duration deviations in auditory stimuli when presented at a medium presentation rate, yet they showed disruptions in their ability to detect duration deviations at slower and faster presentation rates, suggesting an early stage breakdown in auditory sensory memory (Brima, Freedman, et al., 2024). Here, we extend this work by showing preliminary evidence of longitudinal differences in the AEPs of individuals with CLN3 disease relative to NT controls. We specifically found age-related differences when examining the N1 and P2 components of the AEP in individuals with CLN3 disease from ages 11 to 25 years relative to NT controls. These amplitude differences were most notable in CLN3 disease when participants were presented with standard tones that had the slowest presentation rate (∼0.5 Hz), which is consistent with differences seen in the MMN of individuals with CLN3 disease. In contrast, the AEP in CLN3 disease participants looked most similar to controls during the fastest stimulus presentation condition, suggesting intact auditory processing when the auditory sensory memory system is less burdened. Further, our preliminary results showed that the N1 and P2 amplitudes were more negative than NT controls in childhood and adolescence and that as individuals with CLN3 disease aged and their disease progressed, the amplitude of their AEPs appeared to be greater than age-matched controls. However, NT controls showed the opposite pattern across development with a significantly increased negativity of the N1 component across increased age, particularly during the longest stimulus condition. These findings suggest a clear difference in the developmental progression of the AEP in individuals with CLN3 disease compared to typical development.

These results also demonstrate value of translational studies. Specifically, similar auditory processing disruptions have been shown in a mouse model of CLN3 disease using near identical EEG methods (Ding et al., 2025). This report provides compelling evidence of sex-specific auditory processing differences in a CLN3 mouse model during early development. Ding et al., found that male CLN3 disease mice showed an initially greater AEP response in early development followed by a decline and subsequent recovery relative to wild type mice. In contrast, female CLN3 disease mice showed a progressive decline in AEPs relative to female wild type mice (Ding et al., 2025). These mouse model findings nicely parallel findings in our younger CLN3 disease participants (Sub #1 and Sub #2) who were male and female, respectively. In the present study, Sub #1 (male) showed AEPs that were initially greater than child controls and more similar to NT controls during adolescence, while Sub #2 (female) did not show the same pattern of recovery seen in the similar-aged male participant. Our sample is too small to draw clear conclusions, yet our findings emphasize the need for translational research examining sex-based differences in longitudinal changes in AEPs.

The current auditory processing findings also demonstrated associations with clinical measures of cognitive functioning including verbal intelligence and working memory abilities as well as a possible association to CLN3 disease symptom severity. Specifically, a more positive N1 amplitude during the longest stimulus presentation conditions was associated with both reduced verbal intelligence and working memory abilities in individuals with CLN3 disease. Additionally, an increased P2 amplitude during the longer/slower stimulus presentation conditions was associated with an increased disease severity score, however, these results should be interpreted with caution as they are significantly limited by the lack of participants at the early and later disease stages. Nonetheless, the trend of elevated P2 amplitudes at later disease stages should be further explored.

Clinical-translational research in CLN3 disease and other rare childhood-onset neurologic diseases has faced many challenges (Jonker et al., 2025; Mink & Vermilion, 2025). One of the biggest challenges is identifying translational outcome measures that are sensitive to both short-and long-term changes across the disease and can be used in both human and animal models to aid in the transition from pre-clinical to clinical trials. This study has demonstrated that similar EEG methods can be used in both participants with CLN3 disease as well as mouse models (Brima, Freedman, et al., 2024; Ding et al., 2025) and that in both samples, the AEP is sensitive to age and disease related changes. Current clinical trials rely primarily on existing outcome measures including clinician report measures such as the UBDRS. While these measures have been established as reliable and valid tools for the assessment of clinical change in CLN3 disease (Kwon et al., 2011; Masten et al., 2020), it will be advantageous to pair them with objective measures that are more sensitive to changes over a shorter time-period that can serve as potential biomarkers of the disease. A recently published open-label study evaluating the use of Miglustat treatment in a small number of patients with CLN3 disease followed for about 4 years showed that use of the drug was associated with a slower rate of physical decline compared to natural history data in CLN3 disease patients (Pietrafusa et al., 2025). However, the authors note that the development of reliable biomarkers useful for monitoring disease progression and treatment outcomes are necessary to make progress in the development of therapeutics.

While there are several limitations to the current study including the limited number of CLN3 disease participants and the lack of longitudinal data from NT control participants, this study serves as an important demonstration of the use of EEG measures longitudinally in CLN3 disease with the potential of serving as an outcome measure in clinical trials.

## DECLARATIONS

### Ethics approval and consent to participate

All aspects of the research conformed to the tenets outlined in the Declaration of Helsinki, with the exception that this study was not preregistered. The institutional review board of the University of Rochester, where the data collection took place, approved this study (STUDY00000920). All participants provided written informed consent/assent or a waiver of assent, when appropriate.

### Consent for publication

Not applicable

### Author contributions

JJF and EGF conceived the study. JJF, EGF and TB designed the original experiment. HRA, EFA and JV recruited and phenotyped the participants and contributed to review/revision of the manuscript. TB, EKB, SN and ERL collected the data. EKB and ERL analyzed the data and created the illustrations and wrote the first draft of the paper, in close collaboration with JJF and EGF. JJF, EGF, HRA, JV, EFA and TB provided substantial editorial input and writing on subsequent drafts. All authors read the final draft and provided critical input.

## Acknowledgments

Thanks to Jamison Seabury and Síle Ní Mhurchú for help with data collection. We are particularly thankful to the families and participants for their selflessness in participating in this research.

## Availability of data and material

The datasets used during the current study are available upon reasonable request to the corresponding authors.

## Abbreviations List

NCL: Neuronal Ceroid Lipofuscinoses
CLN3: Ceroid Lipofuscinosis Neuronal 3
UBDRS: Unified Batten Disease Rating Scale
CLN3SS: CLN3 Staging System
ERP: Event Related Potential
EEG: Electroencephalography
MMN: Mismatch negativity
AEP: Auditory evoked potential
NT: Neurotypical
WISC-V: Wechsler Intelligence Scale for Children, Fifth Edition
WAIS-IV: Wechsler Adult Intelligence Scale-Fourth Edition
BDSRA: Batten Disease Support and Research Association
VCI: Verbal comprehension index
DSF: Digit span forward
SOA: Stimulus onset asynchrony

## Clinical Trial Number

not applicable.

---

**Supplemental Table 1.**
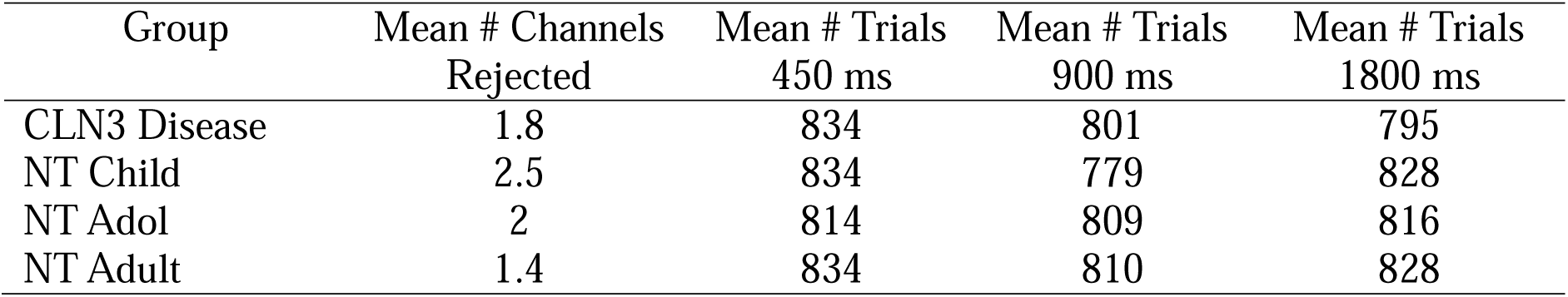
Mean number of channels rejected and trials included for each group and SOA condition.

| Group | Mean # Channels<br>Rejected | Mean # Trials<br>450 ms | Mean # Trials<br>900 ms | Mean # Trials<br>1800 ms |
| --- | --- | --- | --- | --- |
| CLN3 Disease | 1.8 | 834 | 801 | 795 |
| NT Child | 2.5 | 834 | 779 | 828 |
| NT Adol | 2 | 814 | 809 | 816 |
| NT Adult | 1.4 | 834 | 810 | 828 |

**Supplemental Table 2.** Spearman correlations between N1 and P2 amplitudes and clinical measures in CLN3 disease participants.

|  | *N1* |  |  | *P2* |  |  |
| --- | --- | --- | --- | --- | --- | --- |
|  | 450 ms | 900 ms | 1800 ms | 450 ms | 900 ms | 1800 ms |
| DSF | -0.025 | -0.566 | -0.675* | -0.360 | -0.444 | -0.573 |
| VCI | -0.297 | -0.605 | -0.713* | -0.439 | -0.458 | -0.621 |
| CLN3SS | -0.016 | 0.474 | 0.555 | 0.403 | 0.650* | 0.607* |

Note. \* $p < 0.01$ , \*\* $p < 0.001$ ; DSF = Digit Span Forward Scaled Score; VCI = Verbal Comprehension Index Standard Score; CLN3SS = CLN3 Staging System.

**Supplemental Table 3.**
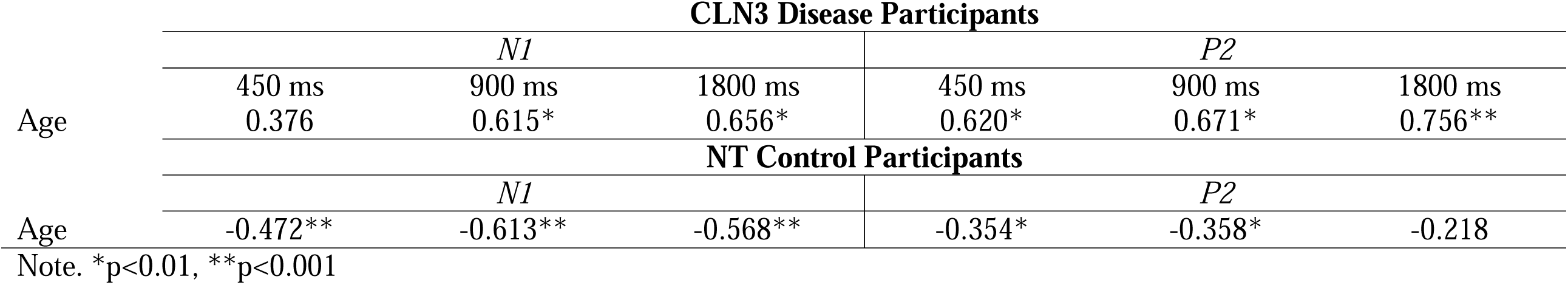
Pearson correlations between N1 and P2 amplitudes and age, separated by CLN3 disease participants and NT control participants.

**Supplemental Figure 1.**
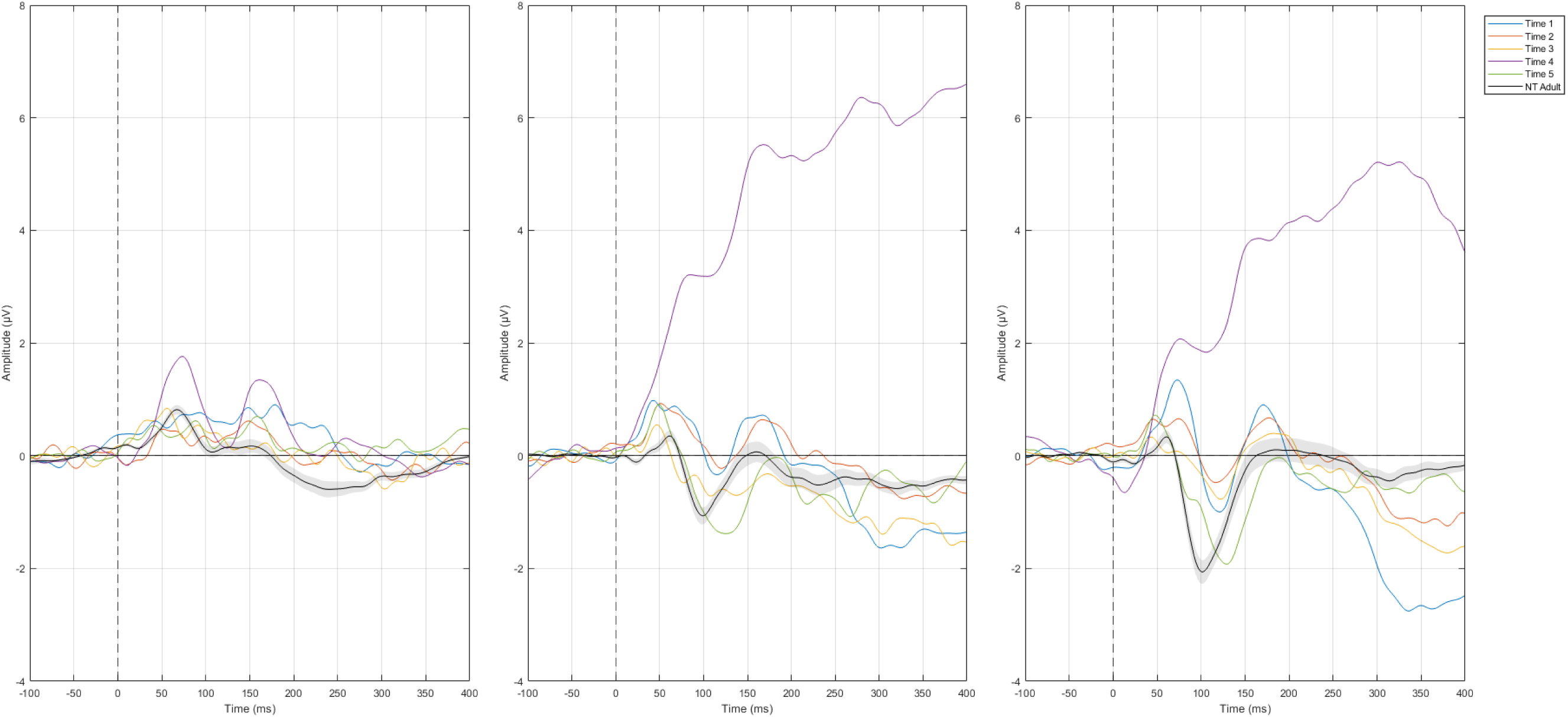
ERPs for CLN3 disease, Subject 3 including timepoint 4 which was excluded from main analyses due to recent major seizure and rescue medication use.

**Supplemental Figure 2.**
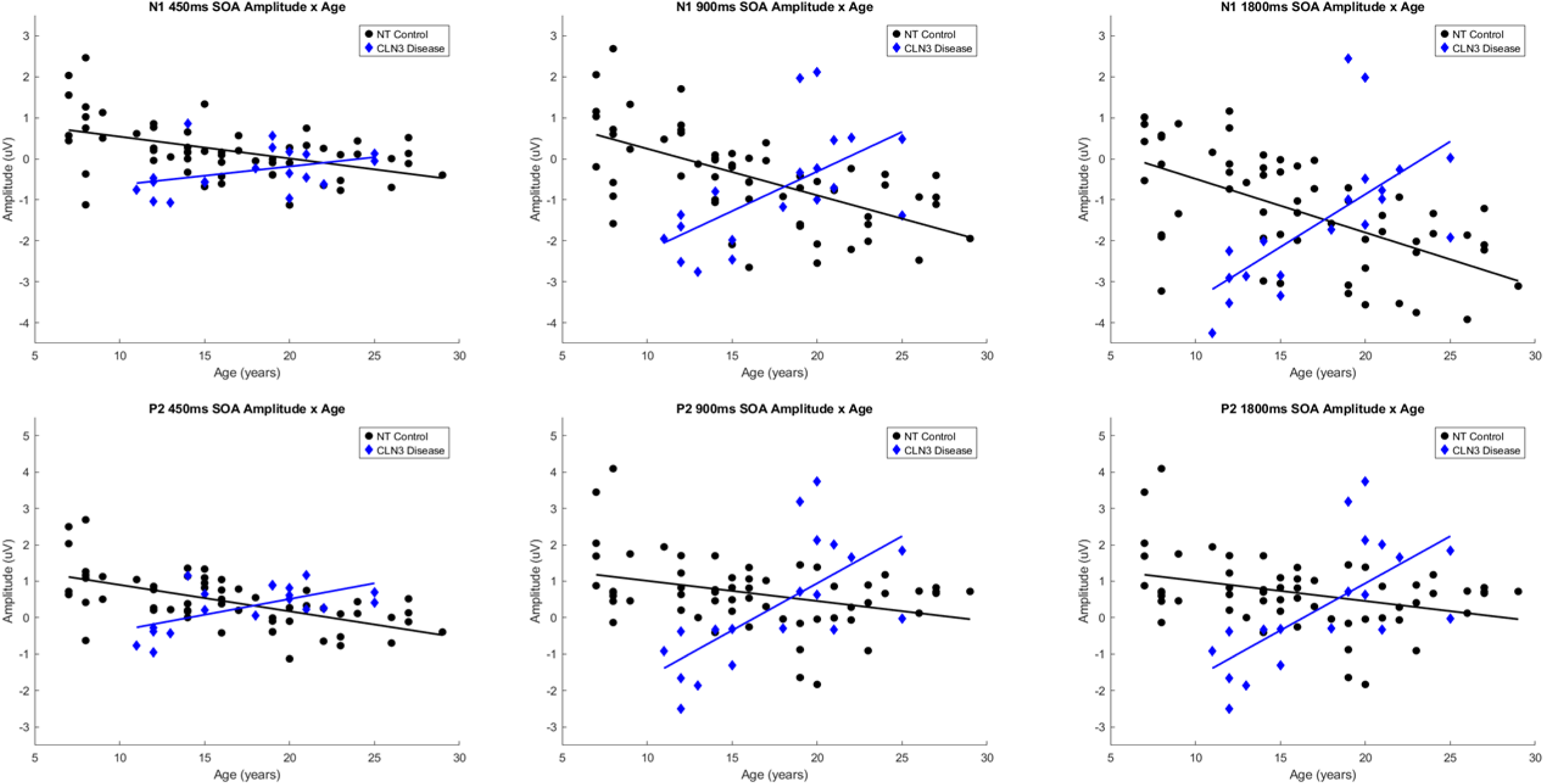
Associations between N1 and P2 amplitudes and age in CLN3 disease and NT control participants.

